# The level of oncogenic Ras controls the malignant transformation of Lkb1 mutant tissue in vivo

**DOI:** 10.1101/2020.09.25.308080

**Authors:** Briana Rackley, Chang-Soo Seong, Evan Kiely, Rebecca E. Parker, Manali Rupji, Bhakti Dwivedi, John M. Heddleston, William Giang, Neil Anthony, Teng-Leong Chew, Melissa Gilbert-Ross

## Abstract

The genetic and metabolic heterogeneity of RAS-driven cancers has confounded therapeutic strategies in the clinic. To address this, rapid and genetically tractable animal models are needed that recapitulate the heterogeneity of RAS-driven cancers in vivo. Here, we generate a *Drosophila melanogaster* model of Ras/Lkb1mutant carcinoma. We show that low-level expression of oncogenic Ras (Ras^Lo^) promotes the survival of Lkb1 mutant tissue, but results in autonomous cell cycle arrest and non-autonomous overgrowth of wild-type tissue. In contrast, high-level expression of oncogenic Ras (Ras^Hi^) transforms Lkb1 mutant tissue resulting in lethal malignant tumors. Using simultaneous multiview light-sheet microcopy, we have characterized invasion phenotypes of *Ras/Lkb1* tumors in living larvae. Our molecular analysis reveals sustained activation of the AMPK pathway in malignant *Ras/Lkb1* tumors, and demonstrate the genetic and pharmacologic dependence of these tumors on CaMK-activated Ampk. We further show that LKB1 mutant human lung adenocarcinoma patients with high levels of oncogenic KRAS exhibit worse overall survival and increased AMPK activation. Our results suggest that high levels of oncogenic KRAS is a driving event in the malignant transformation of LKB1 mutant tissue, and uncover a novel vulnerability that may be used to target this aggressive genetic subset of RAS-driven tumors.

**One Sentence Summary:** A multivariable Ras-driven *Drosophila* model reveals a novel LKB1 mutant lung adenocarcinoma patient subpopulation and targetable effector pathway.

## Introduction

KRAS is the most commonly mutated oncogene in human cancer, and is frequently mutated in cancer types associated with high mortality such as non-small cell lung cancer (NSCLC). Efforts to directly target the KRAS protein have been challenging, although renewed efforts are currently in clinical trials^1^. Large-scale sequencing of lung adenocarcinoma has uncovered heterogeneity in mutant KRAS tumors due to concomitantly mutated tumor suppressor genes such as *TP53* and *LKB1*, genetic subtypes that are largely mutually exclusive and which harbor distinct biologies and therapeutic susceptibilities^2^. An added layer of complexity arises due to the extensive metabolic rewiring observed in RAS-driven tumors^3^, which can arise due to Kras-mutant dosage and alterations in signaling pathways downstream of mutated tumor suppressor genes^4^. Increasingly, metabolic rewiring is known to be dependent on tissue-level dynamics within the tumor and the tumor microenvironment. Therefore, there is a need to develop rapid and powerful models of RAS-driven cancers that mimic the complex landscape of these tumors in vivo.

Liver Kinase B1 (LKB1) is a master serine/threonine kinase that phosphorylates 13 downstream kinases of the AMP-activated protein kinase family (AMPK) family to control cell growth and cell polarity^5^. LKB1 activity is lost in a wide spectrum of human cancers and the gene that encodes LKB1 (*STK11*) is the third most frequently mutated tumor suppressor in human lung adenocarcinoma. Loss of *LKB1* frequently occurs in KRAS-driven lung adenocarcinoma, and has been shown to promote metastasis, shorten overall survival, and confer resistance to targeted therapies and checkpoint inhibitors^6-10^. Altogether, these differences in survival and treatment outcomes highlight the importance of in vivo models that recapitulate the complexity and heterogeneity of these tumors when developing and implementing cancer treatments.

*Drosophila melanogaster* is a powerful model system for studying cancer biology due to the high conservation of human oncogene and tumor suppressor pathways^11,12^. Elegant genetic mosaic techniques in *Drosophila* allow tissue-specific overexpression of oncogenes and knockdown of tumor suppressors within distinct subpopulations of cells, which bestows the ability to build complex tumor landscapes in vivo. Seminal work using these methods has identified cooperating mutations that promote the metastasis of benign *Kras*-mutant tumors in vivo, and has identified such cooperating models as amenable to pharmacologic approaches ^13,14,15,16^. However, despite evidence from mouse models that loss of *Lkb1* is sufficient to promote tumor progression and metastasis in *Kras*-mutant lung tumors^17^, there has been no report of malignant synergy between *Ras* and *Lkb1* using the rapid and genetically tractable *Drosophila* model.

Here, using a novel *Drosophila* model of *Ras/Lkb1*-driven malignant progression, we found that the relative levels of oncogenic Ras determine clonal growth dynamics in *Lkb1* mutant tissue. Low levels of oncogenic Ras promote non-autonomous growth of surrounding wild-type tissue, while high-levels promote malignant progression and organismal lethality. To further characterize the metastatic capability of *Ras/Lkb1* malignant cells we used simultaneous multiview light sheet microscopy to image live tumor-bearing larvae for up to 48hrs, and show that *Ras/Lkb1* cells actively degrade basement membrane, and ultimately invade distant tissues. To further define the mechanism driving the progressive synergy between high oncogenic Ras and loss of *Lkb1* we investigated signaling networks in mosaic tissue. We show that malignant *Ras/Lkb1* tumors activate AMPK and are dependent on the activation of the Drosophila ortholog of CAMKK2. We validate the translational potential of our work by showing high level KRAS with concurrent mutation in *LKB1* represents a unique subset of patients with worse overall survival and increased AMPK activation. Our work uncovers a novel mechanism that may include oncogenic *KRAS* copy number gains or amplification as a novel synergistic mechanism that drives the aggressive nature of *LKB1* mutant tumors. In addition, our work proves *Drosophila* as a powerful model for the rational design of targeted therapies for genetic subsets of RAS-driven cancers, and suggests that the *LKB1* subset of KRAS-driven cancers may benefit from targeting of the CAMKK/AMPK circuit.

## Results

### Clonal loss of Lkb1 in vivo results in autonomous cell death

Recent work has highlighted effects of the dosage of oncogenic Ras on the progression of Ras-dependent cancers^18,19^. Previous work in *Drosophila* has identified myriad pathways that collaborate with mutant Ras to promote tumor progression and metastasis^20^, but how the dosage of Ras affects tumor progression in these multiple hit models is unknown. To address this question, we identified oncogenic Ras transgenes with differing expression levels. One expresses oncogenic Ras at levels similar to endogenous Ras (Ras^Lo^). The other expresses Ras at levels several fold higher (Ras^Hi^) (Fig. 1b). To mimic the genetic landscape of human KRAS-driven cancers we chose to co-mutate the tumor suppressor *LKB1* in Ras^Lo^ and Ras^Hi^ tissue. Most tumor specific *LKB1* mutations are homozygous deletions or loss-of-heterozygosity with somatic mutation^21-23^. Among the latter, nonsense or frameshift mutations leading to protein truncation are the most common^24^. To identify the *Drosophila Lkb1* loss-of-function allele with the strongest reduction in Lkb1 protein levels we first generated an antibody to *Drosophila* Lkb1. We then assayed for Lkb1 protein in transheterozygous larvae using three previously published *Lkb1* loss-of-function alleles (X5^25^), 4B1-11 and 4A4-2^26^) over a large deletion that removes the *Lkb1* gene. The *Lkb1*^X5^ and *Lkb1*^4B1-11^ loss-of-function alleles reduced Lkb1 protein expression by 60% compared to control. However, the *Lkb1*^4A4-2^ allele reduced protein expression by 80% (Fig. 1a), which agrees with prior published genetic data suggesting *Lkb1*^4B1-11^ as having residual protein activity^27^. The *Lkb1*^4A4-2^ allele was chosen for further study and will be referred to as *Lkb1*^-/-^.

**Figure 1.**
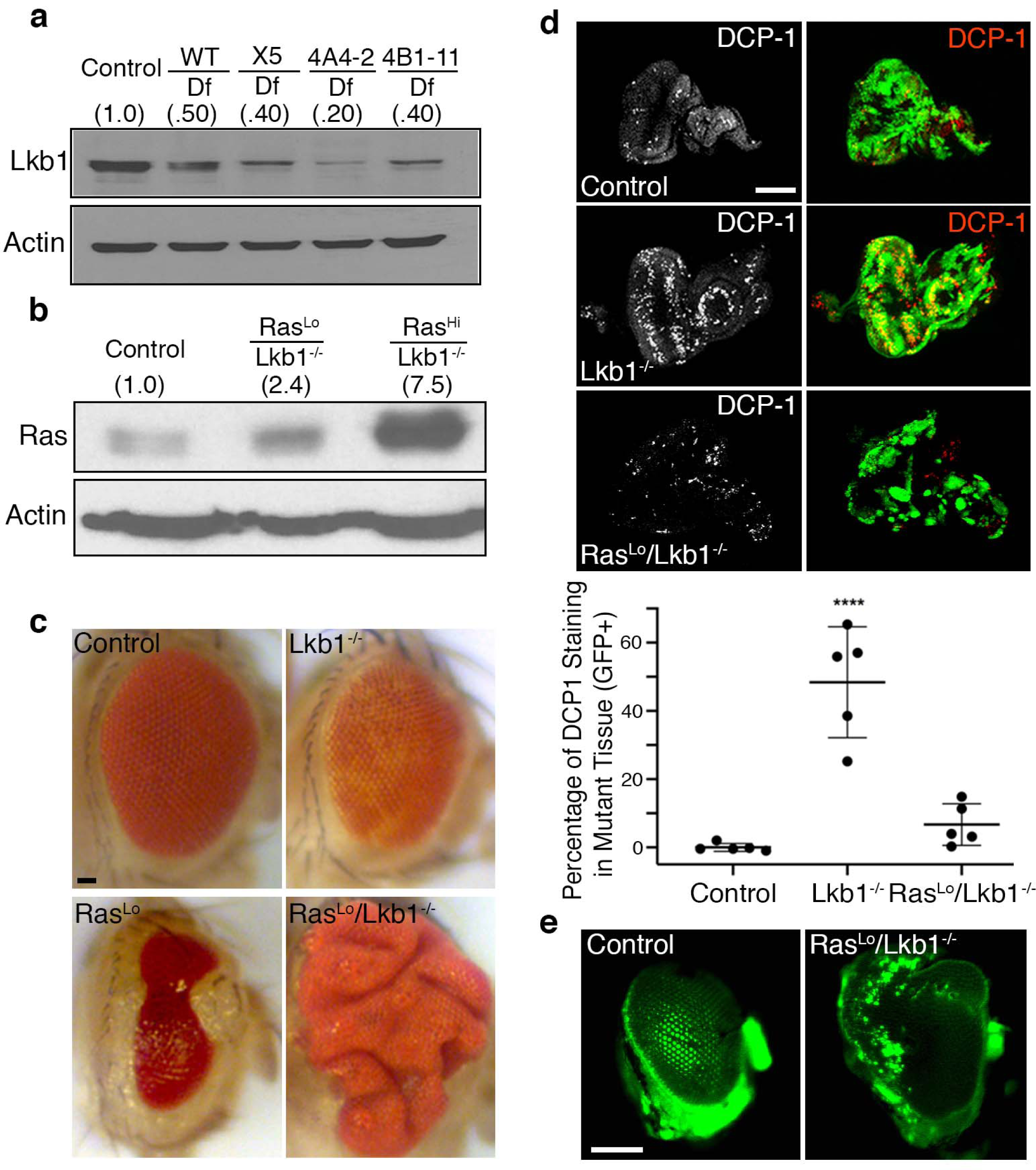
Clonal loss of *Lkb1* in vivo results in autonomous cell death. (a) Western analysis of Lkb1 protein in transheterozygous larvae for a deletion (*Df(3R)Exel6169*) that removes the *Lkb1* gene, and either a wild-type third chromosome or three loss-of-function alleles of *Lkb1* (X5, 4A4-2, 4B1-11). (b) Western analysis of Ras levels in mosaic eye imaginal discs from the indicated genotypes (control = *FRT82B*). Note the Ras antibody detects both endogenous and oncogenic Ras. (c) Representative brightfield images of mosaic adult eyes with clones of the indicated genotypes. Scale bar, 20µm. (d) (top) Confocal maximum intensity projections of third instar mosaic eye discs carrying GFP-tagged clones of the indicated genotypes, and stained for endogenous death caspase 1 (DCP-1). Scale bar, 100µm. (bottom) Percentage of DCP1 staining in GFP-positive mutant tissue was quantified from n=5 imaginal discs per condition using thresholding in FIJI (ImageJ). Data were collected as means +/- SD and plotted using Prism GraphPad. (****P<0.0001, one-way ANOVA with multiple comparisons). (e) Fluorescent images of adult eyes carrying GFP-labelled clones of the indicated genotypes. Images are representative of n=10 independent flies per genotype. Scale bar, 100µm.

We used the GFP-labeled eye expression system^14^ to express Ras^Lo^ in discreet patches or ‘clones’ of developing eye epithelial tissue. Expression of Ras^Lo^ resulted in ablation of eye tissue and benign outgrowths of eye cuticle similar to what has been reported in prior reports using a UAS-Ras^V12^ transgene^14,28^ (Fig. 1c). We then used the GFP-labeled eye expression system to inactivate the *Lkb1* tumor suppressor (*Lkb1*^-/-^) in clones of cells in the developing eye. Inactivation of *Lkb1* in clones resulted in adult flies with small, rough eyes (Fig. 1c), suggesting high levels of apoptosis. To test this, we assayed for cleaved death caspase 1 (DCP-1) in mutant clones using immunofluorescence in wandering 3^rd^ instar eye imaginal discs. As expected, loss of *Lkb1* (marked by GFP+ tissue) resulted in a large increase in autonomous cleaved DCP-1 expression as compared to discs carrying control FRT82B clones (Fig. 1d). These data suggest that homozygous loss of *Lkb1* within an otherwise wild-type epithelium can result in a high level of apoptosis in vivo.

### Low-level Ras and loss of Lkb1 synergize to promote non-autonomous benign overgrowth

Data from genetically engineered mouse models (GEMMs) suggests loss of *Lkb1* is sufficient to promote the progression and metastasis of nascent *Kras* mutant lung adenocarcinoma^17^. Due to the redundancy of the vertebrate genome and paucity of rapid genetic mosaic analyses in GEMMs, we sought to use the GFP-labeled *Drosophila* eye expression system to build a *Ras/Lkb1* model of cooperative tumorigenesis. We simultaneously expressed Ras^Lo^ and depleted *Lkb1* (*Ras*^*Lo*^*/Lkb1*^*-/-*^) in clones of developing eye epithelial tissue, and found that autonomous DCP-1 levels returned to those observed in control eye imaginal discs (Fig. 1d). These data suggest that low levels of oncogenic Ras promote the survival of *Lkb1*^-/-^ mutant tissue in vivo. In addition, eye imaginal disc complexes carrying *Ras*^*Lo*^*/Lkb1*^*-/-*^ clones were larger than mosaic control discs but contained only a small amount of mutant GFP+ tissue compared to the expression of Ras^Lo^ alone. In agreement with these results, analysis of adult *Ras*^*Lo*^*/Lkb1*^*-/-*^ mosaic eyes revealed a large, overgrown eye phenotype composed of mostly GFP-wild-type cells (Fig. 1c,e). To confirm the overgrown eye phenotype was due to synergy between *Ras* and *Lkb1* and not to simply preventing cell death in *Lkb1* mutant cells we expressed the baculoviral caspase inhibitor p35 in *Lkb1* mutant clones. Expressing p35 in *Lkb1* mutant clones resulted in a majority of flies with eyes that are phenotypically similar to expression of p35 alone (normal size eye), with 20% of flies exhibiting a more severe smaller malformed eye (Supplementary Fig. 1).

To investigate the mechanism that results in an increase in organ size in *Ras*^*Lo*^*/Lkb1*^*-/-*^ flies, we analyzed BrdU incorporation in mosaic *Ras*^*Lo*^*/Lkb1*^*-/-*^ eye imaginal disc tissue. Eye disc tissue carrying *Ras*^*Lo*^*/Lkb1*^*-/-*^ clones exhibits BrdU incorporation in GFP-wild-type cells surrounding mutant clones (Fig. 2a). In addition, we analyzed mosaic *Ras*^*Lo*^*/Lkb1*^*-/-*^ eye imaginal disc tissue by fluorescence activated cell sorting (FACS). This analysis revealed an increase in the percentage of GFP+ mutant cells in G1 when compared to GFP+ cells from control FRT82B discs (Fig. 2b,c). Altogether, these data suggest that although *Ras*^*Lo*^*/Lkb1*^*-/-*^ mutant cells survive, they undergo G1 arrest while promoting the increased hyperplastic proliferation of surrounding wild-type tissue.

**Figure 2.**
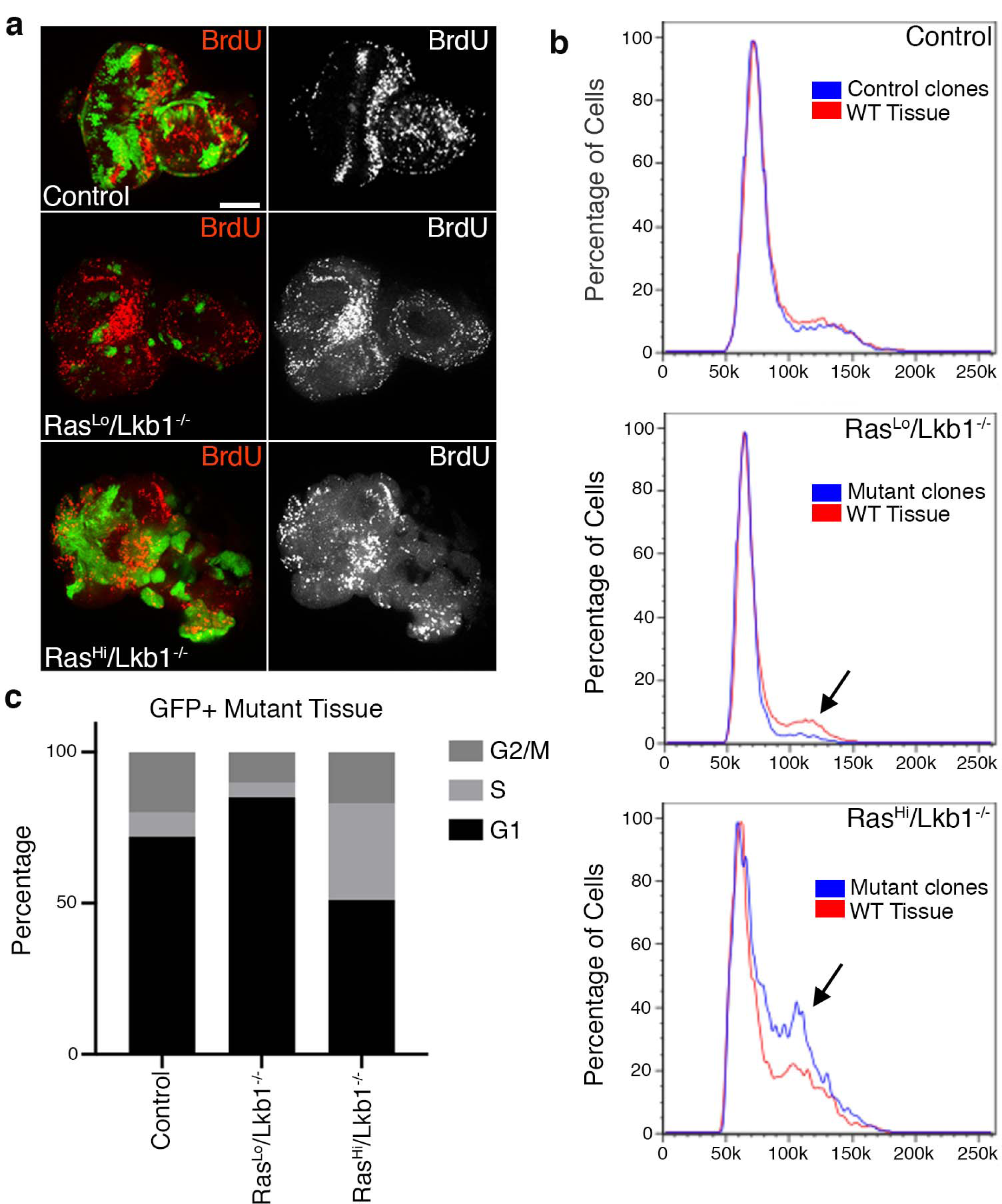
The level of oncogenic *Ras* determines distinct autonomous vs. non-autonomous cell-cycle phenotypes in *Lkb1* mutant tissue. (a) Confocal images of mosaic eye imaginal discs carrying GFP+ clones of the indicated genotypes (control = *FRT82B*), and stained for BrdU incorporation. Images are representative of n=10 independent eye-imaginal discs per genotype. Scale bar, 100µm. (b) Fluorescence-activated cell sorting (FACS) analysis of mosaic eye imaginal discs with GFP-labelled clones of the indicated genotypes. Black arrows point to shifts in relative cell cycle phasing. Analysis is representative of n=3 independent experiments of 20-40 imaginal discs/genotype. c) Histogram showing percentage of GFP-labelled control or mutant cells in each phase of the cell cycle.

### High level oncogenic Kras promotes the neoplastic transformation of Lkb1 mutant tissue

Previous studies have implicated the dose of mutant Kras in tumor progression, cell motility, and metabolic reprogramming^7,18,19,29^, therefore we used the GFP-labeled eye expression system to clonally express *Ras*^*Hi*^ and mutate *Lkb1* in developing eye epithelia (*Ras*^*Hi*^*/Lkb1*^*-/-*^). When combined with *Lkb1* loss-of-function, expression of Ras^Hi^ resulted in severely overgrown and disorganized 3^rd^ instar larval eye-imaginal disc tumors composed of mostly GFP+ mutant tissue (Fig. 3a). FACS analysis of mutant tissue revealed a shift in cell cycle phasing that favored G2/M, suggesting that mutant cells were precociously completing G1 (Fig. 2b,c). The majority of larvae carrying *Ras*^*Hi*^*/Lkb1*^*-/-*^ mosaic discs did not pupate but continued to grow into ‘giant larvae’ while expression of Ras^Hi^ alone resulted in late pupal lethality (Fig. 3b). The giant larval phenotype is shared by loss-of-function mutations in the *Drosophila* neoplastic tumor suppressor genes^30^ and suggests that *Ras*^*Hi*^*/Lkb1*^*-/-*^ tumors are malignant. To test this, we performed an allograft assay by implanting control, *Ras*^*Lo*^*/Lkb1*^*-/-*^, and *Ras*^*Hi*^*/Lkb1*^*-/-*^ GFP+ tumor tissue in the abdomens of wild-type hosts. Transplanted control and *Ras*^*Lo*^*/Lkb1*^*-/-*^ tissue failed to grow in host abdomens (Fig. 3e). Surprisingly, the lifespan of hosts with transplanted *Ras*^*Lo*^*/Lkb1*^*-/-*^ tissue was shortened which suggests that residual GFP-‘wild-type’ tissue from the transplant could be partially transformed. In contrast, only transplanted *Ras*^*Hi*^*/Lkb1*^*-/-*^ tissue was able to grow into visible secondary tumors that significantly shortened host survival (Fig. 3e,f) thus confirming the malignancy of *Ras*^*Hi*^*/Lkb1*^*-/-*^ tumor tissue.

**Figure 3.**
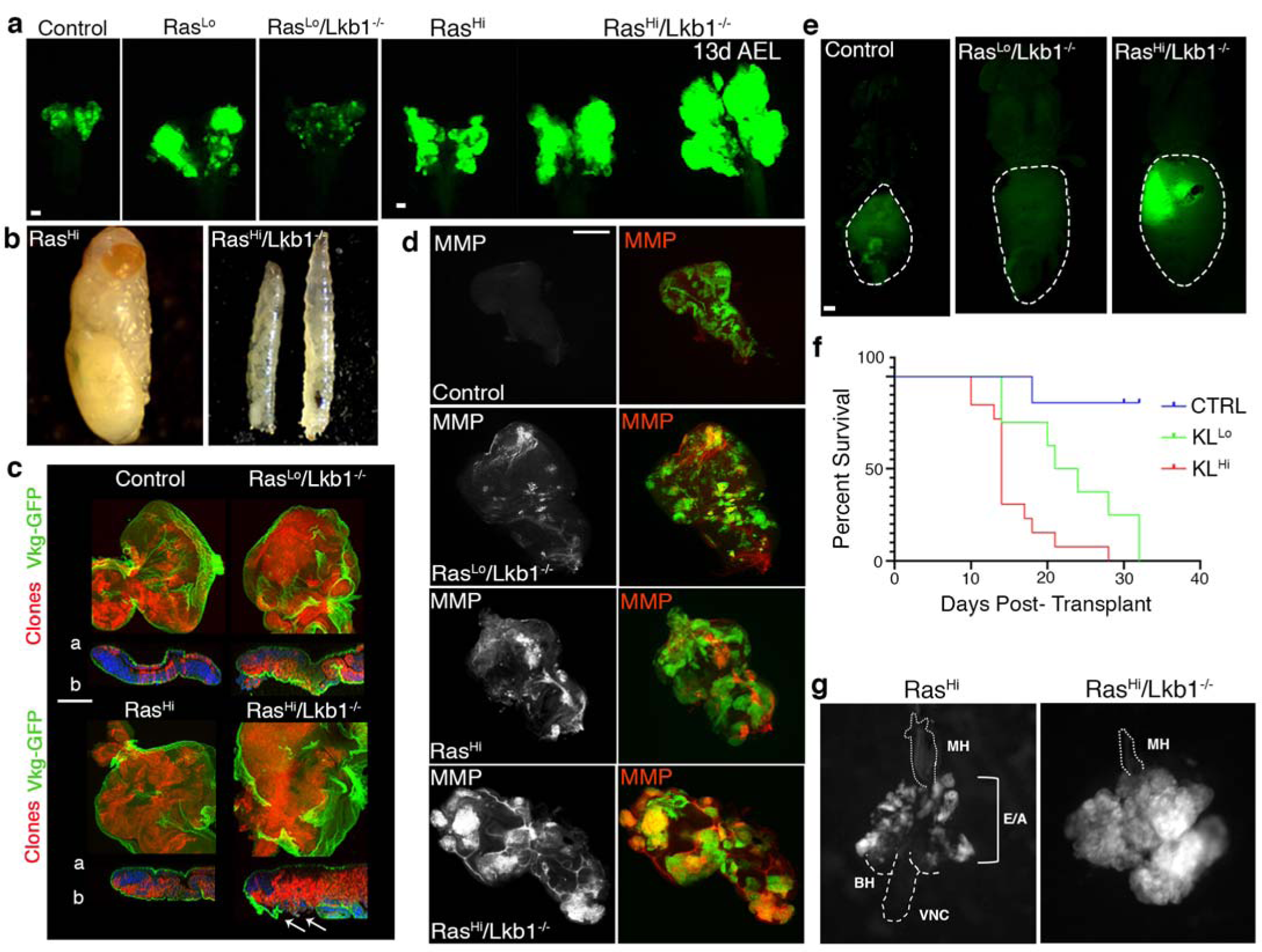
Oncogenic *Ras*^*Hi*^ promotes the malignant transformation of *Lkb1* mutant tissue. (a) Fluorescent images of 3^rd^ instar larval eye-imaginal discs (exception labeled ‘13d AEL’) carrying GFP^+^ clones of the indicated genotypes (control = *FRT82B*). Images are representative of n=10 independent eye-imaginal discs per genotype. Scale bar, 20µm. AEL = after egg lay. The stage ‘13 days AEL’ is indicative of a larva that failed to pupate at day ∼5d AEL, and is a classic neoplastic phenotype. (b) Representative brightfield image of the lethal stage of a fly carrying Ras^Hi^ clones (left) and *Ras*^*Hi*^*/Lkb1*^*-/-*^ clones (right). Note that both age-matched third instar and giant larvae are shown for the *Ras*^*Hi*^*/Lkb1*^-/-^ genotype. (c) Confocal images of eye imaginal discs carrying RFP^+^ clones of the indicated genotypes and expressing type IV collagen-GFP (Vkg-GFP). Nuclei are labelled with DAPI (blue). White arrow indicates breaks in Vkg-GFP. a= apical, b= basal. Images are representative of n=5 independent eye-imaginal discs per genotype. Scale bar, 100µm. (d) Confocal images of third instar eye discs carrying GFP^+^ clones of the indicated genotypes, and stained for matrix metalloproteinase 1 (MMP1). Images are representative of n=10 independent eye-imaginal discs per genotype. Scale bar, 100µm. (e) Fluorescent images of w^1118^ adult virgin female hosts carrying transplanted allografts of 3^rd^ instar eye-imaginal discs with GFP^+^ clones of the indicated genotypes. Images are representative of n=5-10 independent hosts per genotype. Scale bar, 100µm. (f) Quantification of survival post-transplant in allograft assay. Survival was measured from n >5 independent adult hosts per genotype and graphed using a Kaplan-Meier survival plot. Survival post-transplant was measured from 7 days post-transplant to time of death. CTRL (FRT82B), KL^Lo^ (*Ras*^*Lo*^*/Lkb1*^*-/-*^), and KL^Hi^ (*Ras*^*Hi*^*/Lkb1*^*-/-*^) (CTRL-KL^Lo^, **P=0.0030, CTRL-KL^Hi^, ***P=0.0001, KL^Lo^-KL^Hi^, *P=0.0129, Log-rank test). (g) Representative fluorescent images of dissected cephalic complexes and ventral nerve cord (VNC) from larvae carrying GFP^+^ clones (white) of the indicated genotypes. BH = brain hemispheres; E/A = eye/antennal discs, MH = mouth hooks. The *Ras*^*Hi*^*/Lkb1*^-/-^ tissue completely invades and obscures contiguous organs. Images are representative of n=10 cephalic complexes/genotype.

### High-level Ras promotes the invasion and metastasis of Lkb1 mutant tissue

Mutations in cell polarity proteins cooperate with oncogenic Ras to drive tumor cell invasion and metastasis^20^. Previous studies have shown that Lkb1 regulates cell polarity and epithelial integrity across species^31,32^, therefore, we hypothesized that malignant *Ras*^*Hi*^*/Lkb1*^*-/-*^ tumors would have invasive properties. To test this, we first examined whether *Ras/Lkb1* mutant cells compromised basement membrane structure by examining the expression of GFP-tagged Collagen IV (Viking (Vkg)-GFP) using conventional fixation and confocal microscopy. Compared to control and *Ras*^*Lo*^*/Lkb1*^*-/-*^ tissue which shows contiguous Vkg-GFP expression in epithelia, *Ras*^*Hi*^*/Lkb1*^*-/-*^ tissue exhibits breaks in Vkg-GFP expression (Fig. 3c). Expressing Ras^Hi^ on its own is lethal (albeit at the pharate adult stage), so we investigated Vkg-GFP in this genotype and once again found no breaks in the structure of Vkg-GFP. We next assayed matrix metalloproteinase (MMP) expression, as MMPs degrade basement membrane. Compared to control, *Ras*^*Lo*^*/Lkb1*^*-/-*^,and *Ras*^*Hi*^ clones, *Ras*^*Hi*^*/Lkb1*^*-/-*^ mutant tissues express high-levels of autonomous MMPs (Fig. 3d). Lastly, we measured the extent to which *Ras*^*Hi*^*/Lkb1*^*-/-*^ cells invade local tissues by dissecting cephalic complexes and assaying extent of migration over the ventral nerve cord (VNC). Compared to Ras^Hi^ control tissue which exhibits benign overgrowths confined to the eye-antennal discs, *Ras*^*Hi*^*/Lkb1*^*-/-*^cells completely invade contiguous organs like the brain hemispheres and VNC (Fig. 3g). These data suggest *Ras*^*Hi*^*/Lkb1*^*-/-*^ tumor cells escape the basement membrane using an active proteolytic process and invade local tissues.

Invasion and metastasis are difficult processes to visualize in living organisms. Thus far, *Drosophila* tumor-bearing larvae have been precluded from fast, high resolution long-term intravital imaging techniques due to their size, degree of movement, and the significant amount of light scattering throughout the body due to the larval cuticle. To address this, we prepared live tumor-bearing larvae for long-term intravital imaging and used simultaneous multiview (SiMView) light-sheet microscopy (see Methods and ^33^) to visualize tumor cell and collagen IV dynamics for up to 48hrs. SiMView allowed imaging of rapid cellular processes over time on an organismal scale, with minimal photobleaching. We collected image stacks in the *z* range every 60-s on two individual ‘giant’ tumor-bearing larvae with RFP-tagged *Ras*^*Hi*^*/Lkb1*^*-/-*^ mutant cells and Vkg-GFP expressed in the basement membrane of all epithelial tissues. Breakdown of Vkg-GFP was visible over time in each individual larva, especially in overlying tracheal branches dorsal to the tumor surface (Fig. 4a and Supplemental Video 1). We defined two independent regions of interest in each larva that encompassed a tumor-adjacent tracheal branch and calculated Vkg-GFP pixel intensity every 2 hours over a 14hr imaging window. Using the wing disc of each animal as an internal control, we observed a statistically significant difference in the change in levels of Vkg-GFP over the imaging window in the tracheal branches (Fig. 4b). Volumetric rendering and surface reconstruction of the tracheal branches revealed tumor cells in contact with trachea at several hundred µm away from the primary tumors (Fig. 4c-e)) and on rare occasion were found on the ‘interior’ surface of Vkg-GFP. These data suggest *Ras*^*Hi*^*/Lkb1*^*-/-*^ mutant cells actively invade tracheal vascular cells to potentially spread to distant organs.

**Figure 4.**
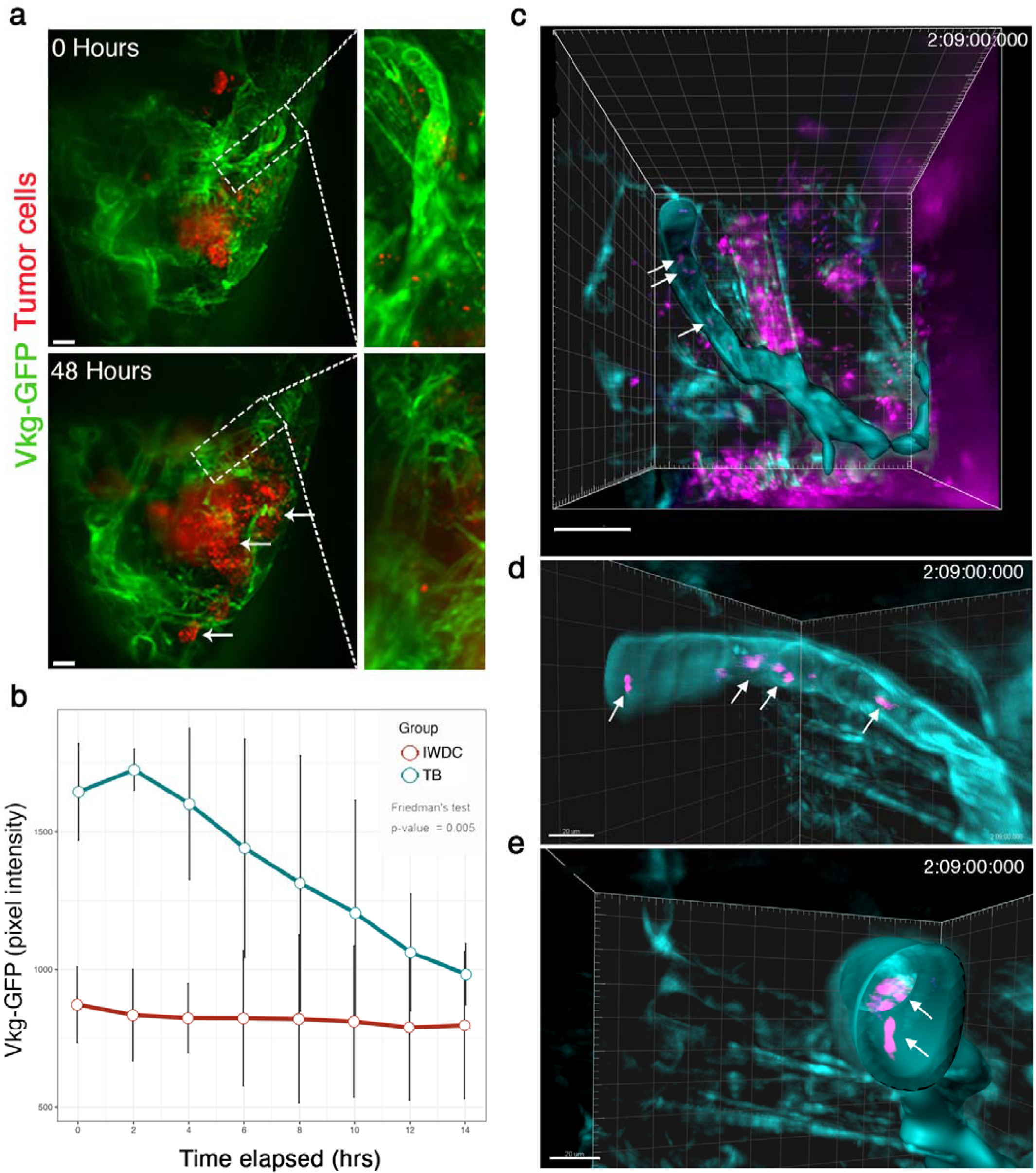
SiMView light sheet microscopy allows visualization of collagen IV degradation by tumor cells over time. (a) Maximum intensity projection (MIP) volume renders from a 48hr SiMView imaging session on the anterior end of a *Ras*^*Hi*^*/Lkb1*^*-/-*^ tumor bearing ‘giant’ larva (13 days AEL). Mutant cells express RFP and Vkg-GFP (collagen IV-GFP) is expressed throughout the organism. White dashed box is a representative region of interest (ROI, tracheal branch) and is magnified in the panels on the right. White arrows indicate RFP-positive *Ras*^*Hi*^*/Lkb1*^*-/-*^ cells that have invaded dorsally. Scale bar, 20µm. (b) The effect of tracheal branch (TB) vs internal wing disc control (IWDC) groups Viking-GFP pixel intensity on time elapsed. Error bars represent standard error of the mean. The group effect was measured using Friedman’s test. (c) An Imaris Surface object of *Vkg-GFP* (teal) was generated from the ROI (above) using min and max thresholds of 250 and 385, respectively. White arrows indicate RFP positive tumor cells (magenta) that appear embedded within the tracheal collagen matrix. (d-e) Zoom and rotated data channels were duplicated with voxels outside the Imaris object set to 0 in order to allow for better visualization with a maximum intensity projection view and clipping plane to show presence of RFP-positive cells within the tracheal matrix. Scale bar, 20µm.

### Ras^Hi^/Lkb1^-/-^ malignant tumors depend on CaMK/Ampk signaling in vivo

Targeting effector signaling in KRAS-driven NSCLC has resulted in limited efficacy in the clinic. In addition, previous studies have highlighted the additional complex transcriptional and signaling network changes in KRAS-driven tumors co-mutated for the tumor suppressor *LKB1*^5^. Therefore, rapid and genetically tractable models of *Kras/Lkb1* tumors may shed light on the complex cring of signaling pathways and highlight novel targeting approaches. To probe effector pathways in our tumor model we used Western analysis on a panel of *Drosophila* epithelia harboring mutant clones for *Ras*^*Lo*^, *Ras*^*Lo*^*/Lkb1*^*-/-*^, *Ras*^*Hi*^, and *Ras*^*Hi*^*/Lkb1*^*-/-*^. Similar to human *KRAS/LKB1* tumors, increases in the activation of the RAS effector circuit Erk/Mek were observed along with S6K and 4EBP1 suggesting increased mTOR pathway activity (Fig. 5a). Compared to all other genotypes AKT is not active in *Ras*^*Hi*^*/Lkb1*^*-/-*^ cells most likely owing to sustained pS6K signaling resulting in a negative feedback loop by ribosomal protein S6. Previous studies have attributed increased TOR pathway activity in *LKB1* mutant tissue to loss of mTOR pathway inhibition by AMPK^36^. Therefore, we tested for loss of AMPK activity in our panel of *Lkb1*^-/-^ mutant *Drosophila* tissue. We observed basal activation of Ampk in control tissue, followed by minimal activation in *Ras*^*Lo*^*/Lkb1*^*-/-*^ mutants, most likely resulting from the overgrowth of surrounding wild-type epithelial tissue (Fig. 5b). However, in *Ras*^*Hi*^*/Lkb1*^*-/-*^ tissue we observed sustained pAmpk levels. Recently, the presumed role of Ampk as a tumor suppressor has been challenged by evidence that Ampk can promote metabolic adaptation to effect tumor growth and survival^38^. To test whether *Ras*^*Hi*^*/Lkb1*^*-/-*^ tumors are dependent on the genetic dose of *ampk* we expressed an RNAi transgene to *ampk* (knockdown efficiency of 50%, see Supplemental Fig. 2a) in developing GFP+ *Ras*^*Hi*^*/Lkb1*^*-/-*^ tissue. Inhibition of Ampk via RNAi in *Ras*^*Hi*^*/Lkb1*^*-/-*^ mutant clones resulted in a statistically significant percentage of flies surviving to adulthood. Interestingly, surviving flies exhibited hyperplastic overgrowth similar to that of Ras^Lo^/Lkb1^-/-^ adult flies (compare Fig. 5d to 1c). A recent study from the Guo group found that autophagy may sustain AMPK activity upon *Lkb1* loss to support tumor growth^39^. In support of this, we detected increased lipidated ATG8a in *Ras*^*Hi*^*/Lkb1*^*-/-*^ tumors, indicative of an increase in autophagic flux (Fig. 5a). Altogher, these data support the conclusion that activation of Ampk is maintained in *Ras*^*Hi*^*/Lkb1*^*-/-*^ tumors and is required autonomously to promote malignant progression of *Kras/Lkb1* tumors in vivo.

**Figure 5.**
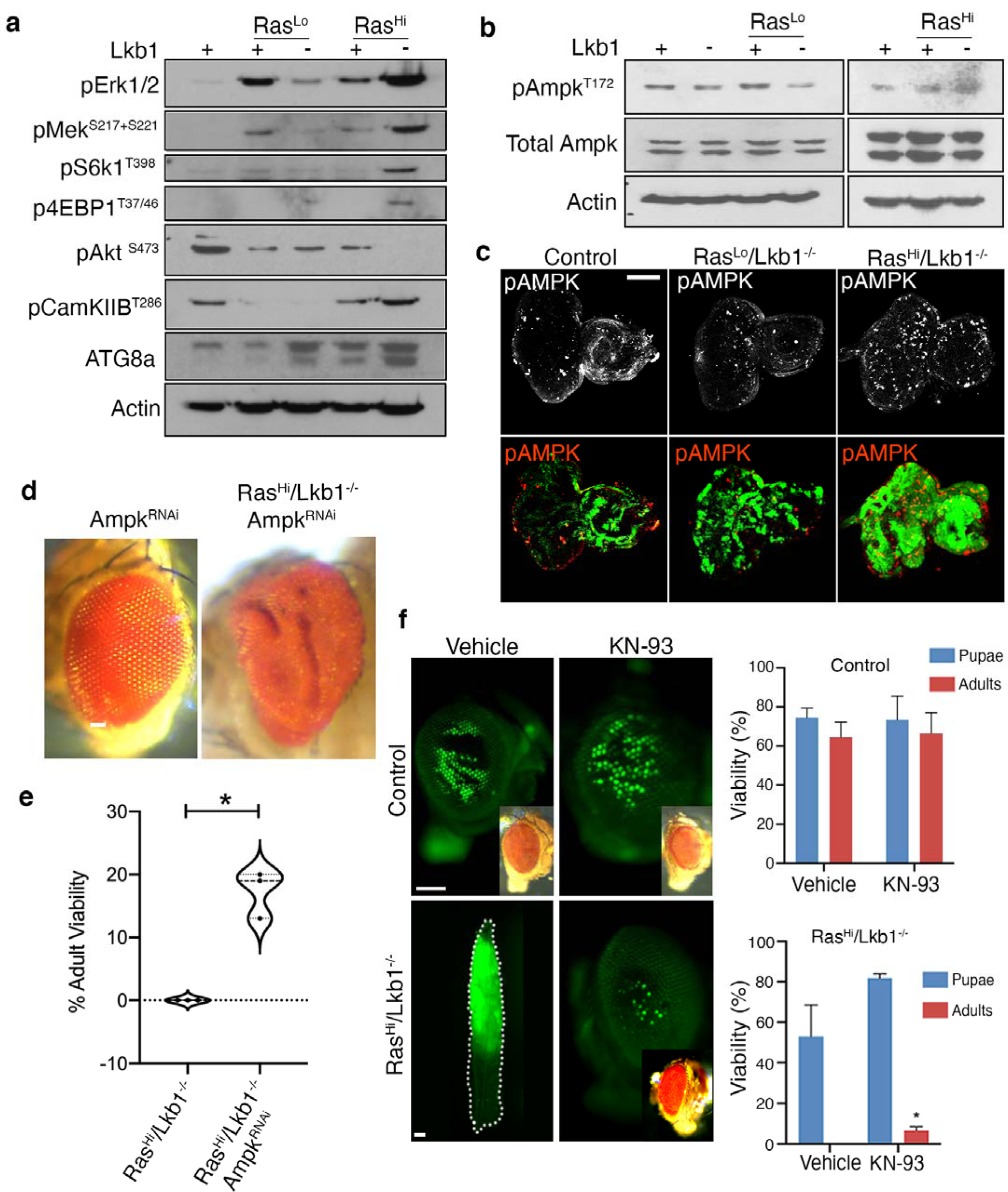
Neoplastic *Ras*^*Hi*^/Lkb1^-/-^ tumors depend on the genetic dose of ampk and are targetable with a CaMK inhibitor. (a) Western analysis to assay activation of the indicated molecular pathways in mosaic larval eye-imaginal discs of the indicated genotypes. (b) Western analysis of Ampk activation from mosaic larval eye imaginal discs of the indicated genotypes. c) Confocal images of eye imaginal discs carrying GFP+ clones of the indicated genotypes (control = *FRT82B*), and stained for phosphorylated Ampk. Images are representative of n=10 independent eye-imaginal discs per genotype. Scale bar, 100µm. (d) Representative brightfield images of adult eyes carrying clones of the indicated genotypes. (e) Violin plot showing % adult viability from the Ras^Hi^/Lkb1^-/-^ (n = 204) and Ras^Hi^/Lkb1^-/-^/Ampk^RNAi^ (n = 139) genotypes. Data is pooled from 3 independent experiments, *p-value = .0.0155 using a Welch’s t-test. (f) Representative fluorescent and brightfield (inset) images of either flies carrying control (*FRT82B*) or GFP^+^ *Ras*^*Hi*^*/Lkb1*^*-/-*^ clones that were pharmacologically treated with vehicle or the pan CaMK inhibiter KN-93 (5µM) as 1^st^ instar larvae. The percent survival to pupal and adult stages was quantified (right). Data were plotted as percentages of total, with two separate experiments for a total of n=50 larvae per condition, *p-value = .0.0493. Scale bars, 100µm.

The Ca2+/calmodulin-dependent protein kinase kinase (CaMKK2) is a nucleotide-independent activator of AMPK^37^, therefore we assayed activation of the *Drosophila* ortholog CamkIIB (48% identical/63% similar to CaMKK2) in our panel of mutant tissue. We found that activation of CamkIIB was elevated in Ras^Hi^/Lkb1^-/-^ tumors (Fig. 5a), suggesting a conserved role for this kinase in activating Ampk in the presence of oncogenic Ras tumors lacking Lkb1. To test whether Ras^Hi^/Lkb1^-/-^ tumors are dependent on CamkIIB activity we used pharmacologic inhibition of the CaMK cascade by feeding developing Ras^Hi^/Lkb1^-/-^ larvae with the inhibitor KN-93^40^, which in our model inhibited activation of the *Drosophila* CamkIIB by 47% (Supplemental Fig. 2b). Treatment of Ras^Hi^/Lkb1^-/-^ larvae resulted in a significant rescue of whole-organismal lethality, with an increase in the number of flies surviving to the pupal and adult stage (6.5% adult survival for KN-93 vs. 0% adult survival for vehicle control) (Fig. 5f). Taken together, these data suggest that in the context of loss of Lkb1, high levels of oncogenic Ras result in activation of Ampk by the alternative sole *Drosophila* CAMKK2 ortholog. Moreover, our pharmacologic results suggest that targeting the upstream AMPK/CAMKK complex may offer therapeutic benefit to *KRAS/LKB1* mutant lung adenocarcinoma patients.

### High levels of oncogenic KRAS and loss of LKB1 result in decreased patient survival and AMPK signaling circuit activation in the TCGA lung adenocarcinoma cohort

To test the translational relevance of our findings in *Drosophila* we analyzed human lung adenocarcinoma genomic and clinical data using cBioPortal^41,42^ to study how differences in levels of oncogenic KRAS affect tumor progression in *LKB1* mutant patients. We used the TCGA Lung Adenocarcinoma PanCancer Atlas and TCGA Provisional Lung Adenocarcinoma datasets to select the proportion of patients with KRAS mutations in codon 12 (G12C, G12D, or G12V) for further study. We then used available RNA sequencing data to stratify patients as either RAS^Lo^ or RAS^Hi^. We next investigated overall patient survival by comparing cohorts of RAS^Lo^ or RAS^Hi^ alone, to those that contained mono, bi-allelic loss and/or loss-of-function mutations in *LKB1*. We found no difference in overall survival in *RAS*^*Lo*^ vs. *RAS*^*Lo*^*/LKB1*^*Mut*^ patients, but strikingly *RAS*^*Hi/*^*LKB1*^*Mut*^ patients exhibited significantly worse overall survival when compared with *RAS*^*Hi*^ patients (Fig. 6a,b). We then tested whether *KRAS* copy number changes could account for the change in overall survival. Similar results were obtained when patients were stratified into either oncogenic *Ras*^*Diploid*^ or *Ras*^*Gain/Amp*^ (Fig. 6c,d). Interestingly, the ability of high level vs. low level KRAS to drive survival differences did not extend to patients with *TP53* mutations (Supplemental Fig. 3).

**Figure 6.**
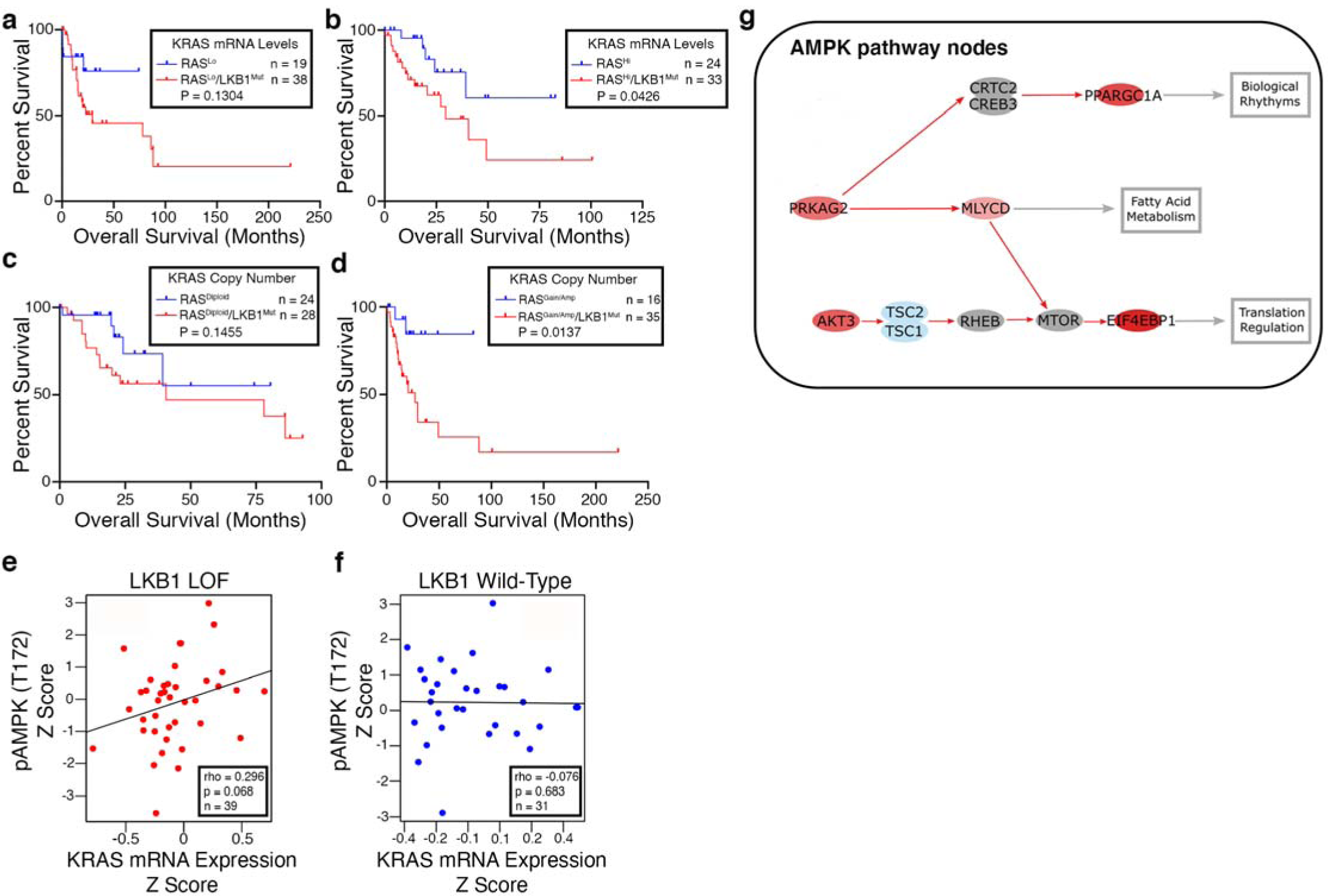
High levels of oncogenic *KRAS* drive decreased patient survival and is associated with AMPK activation in *LKB1* mutant patients. (a-b) Analysis of patient survival using the TCGA Pan Lung Cancer study. Kaplan Meier plots stratified by RAS^Lo^ or RAS^Hi^ using oncogenic (codon 12) *KRAS* mRNA expression and further stratified based on *LKB1* deletion and loss-of-function mutation status. (c-d) Analysis of patient survival using the TCGA Pan Lung Cancer study. Kaplan Meier plots stratified by RAS^Lo^ or RAS^Hi^ using oncogenic (codon 12) *KRAS* copy number data and further stratified based on *LKB1* deletion and loss-of-function mutation status. (e-f) Analysis of phosphorylated AMPK (T172) expression as it correlates with *KRAS* mRNA expression and *LKB1* mutation status. (g) Canonical circuit activity analysis (CCAA) (http://hipathia.babelomics.org) was used to estimate the activity of AMPK signaling pathway (hsa04152) that result in functional cell activities. Red color represents significantly (p<0.05) upregulated genes (or paths) in *KRAS*^*Hi*^*/LKB1*^*Mut*^ lung adenocarcinoma patients with respect to *KRAS*^*Hi*^ patients, and blue represents downregulated genes (or paths). The activity of three effector circuits are significantly (FDR<0.05) upregulated in *KRAS*^*Hi*^*/LKB1*^*Mut*^ patients, one ending in the node that contains the protein PPARGC1A (p = 0.005; FDR = 0.037; Uniprot function Biological rhythms/Mitochondrial biogenesis), the second one ending in the node with the MLYCD protein (p = 0.0064; FDR = 0.045; Uniprot function Fatty acid metabolism), and the third ending in the node containing EIF4EBP1 (p = 0.001; FDR = 0.013; Uniprot function Translation regulation).

A recent study has reported that AMPK has a pro-tumorigenic role in lung cancer GEMMs with Kras and p53 mutations^43^. Moreover, data from our *Drosophila Lkb1* mutant tumor model indicate that halving the genetic dose of *ampk* is sufficient to partially reverse whole-organism lethality. To test whether AMPK signaling may be involved in human *KRAS/LKB1* mutant lung adenocarcinoma we performed a correlation analysis between pAMPK and oncogenic codon 12 *KRAS* mRNA for *LKB1* loss-of-function and *LKB1* wild-type patients using TCGA data. We detected a positive correlation trend between pAMPK and oncogenic *KRAS* levels, but only in *LKB1* mutant patients (Spearman’s correlation coefficient = 0.3, p = 0.068 for LKB1 loss-of-function vs coefficient = −0.076, p = 0.683 for LKB1 wild-type patients) (Fig. 6e,f). To further test our hypothesis, we used canonical circuit activity analysis (CCAA)^44^ which recodes gene expression data into measurements of changes in the activity of signaling circuits, ultimately providing high-throughput estimations of cell function. We performed CCAA to estimate activity of the AMPK pathway in *RAS*^*Hi*^*/LKB1*^*Mut*^ lung adenocarcinoma patients compared to *RAS*^*Hi*^ patients. The activity of three effector circuits are significantly (FDR<0.05) upregulated in *KRAS*^*Hi*^*/LKB1*^*Mut*^ patients, one ending in the node that contains PPARGC1A (encodes PGC1alpha), the second one ending in the node with the MLYCD gene, and the third ending in the node containing EIF4EBP1 (Fig. 6g). These three genes control the cellular processes of circadian control of mitochondrial biogenesis, fatty acid metabolism, and translation regulation, and are known to be upregulated in various cancers^45,46,47^. These data confirm the translational relevance of our *Drosophila* model, and suggest that high oncogenic *KRAS* levels, perhaps through copy number gains, activate specific sub-circuits of the AMPK signaling pathway to drive the malignant progression of *LKB1* mutant tumors.

## Discussion

Co-occurring genomic alterations in oncogene-driven lung adenocarcinoma are emerging as critical determinants of tumor-autonomous and non-autonomous phenotypes^2^. Here, we have generated the first *Drosophila* model of *Ras/Lkb1* co-mutation, a major subgroup of *KRAS*-driven lung adenocarcinomas. Our results indicate that the levels of oncogenic *Ras* determine key autonomous vs. non-autonomous phenotypes in *Lkb1* mutant tissue. Low-level oncogenic Ras expression (Ras^Lo^) combined with *Lkb1* co-mutation results in autonomous G1 arrest and overgrowth of the surrounding wild-type epithelium. Conversely, high-level oncogenic Ras (Ras^Hi^) combined with *Lkb1* co-mutation leads to autonomous transformation, invasion, and metastasis.

It has been proposed that RAS-induced senescence functions as a tumor suppressive mechanism^48^. More recent data have built upon these studies to show that high levels of Hras are required to activate tumor suppressor pathways in vivo^18^, and that doubling the levels of oncogenic *Kras* is sufficient to cause metabolic rewiring leading to differences in therapeutic susceptibilities^19^. Mutant *Kras* copy gains are positively selected for during tumor progression in a *p53* mutant background^49^; however, our results analyzing survival in patients indicate that unlike *KRAS/LKB1*, high levels of *KRAS* in *TP53*-mutant lung adenocarcinoma patients may not be a key factor in determining overall survival. In contrast, high-level *KRAS* and loss of *LKB1* leads to significantly decreased overall survival in lung cancer. Interestingly, *LKB1* has been shown to control genome integrity downstream of DNA damaging agents and cellular accumulation of ROS. Moreover, alterations in *LKB1* occur more frequently in patients with no known mitogenic driver^50^. Future work should uncover whether *KRAS* copy number gains and amplifications are positively selected for due to the role of LKB1 as a gatekeeper of genome integrity.

Seminal work in *Drosophila* identified the loss of epithelial polarity genes as key cooperating events in Ras-driven tumors in vivo^13,14^. In addition to its role in regulating cell growth, the Lkb1 protein is required to establish and maintain cell polarity across eukaryotes. However, alleles of *Lkb1* were not reported to synergize with oncogenic Ras in these studies, the reason possibly due to insufficient oncogenic Ras levels. The fact that loss of *Lkb1* behaves differently than other known polarity mutants suggests that an alternate function underlies the aggressive nature of *Lkb1* mutant cancer. Our work for the first time shows that functional Ampk activity is required for the malignant progression of *Ras*^*Hi*^*/Lkb1*^*-/-*^ tumors in vivo. Moreover, our findings in lung adenocarcinoma patients suggest that increased oncogenic *KRAS* is associated with increased activation of pAMPK in *LKB1* mutant patients. The fact that pharmacologic inhibition of the *Drosophila* Camkk2 ortholog using the compound KN-93 resulted in partial suppression of Kras^Hi^/Lkb1^-/-^ larval/pupal lethality further suggests that high-level oncogenic signaling engages the CaMK pathway to activate Ampk in *Ras/Lkb1* mutant tissue.

Using CCAA, we discovered PPARGC1A, which encodes the protein PGC1α, as significantly upregulated in KRAS^Hi^/LKB1^Mut^ lung adenocarcinoma patients. Interestingly, studies in human prostate cancer have discovered metabolic adaptations through PGC1α-mediated mitochondrial biogenesis in response to CAMKKβ/AMPK signaling^51,52^. Future studies should focus on whether similar adaptations drive tumor growth and survival in *KRAS/LKB1* mutant lung adenocarcinoma. In addition, work is needed to elucidate the mechanism used by high-level Ras signaling to engage the CaMK pathway. Lastly, our work is the first to show that Ampk can have a pro-tumorigenic role in *Lkb1* mutant cancer in vivo, and suggests that *KRAS/LKB1* mutant lung adenocarcinoma patients may benefit from CAMKK inhibitors.

## Materials and Methods

### *Drosophila* stocks and maintenance

Flies were grown on a molasses-based food at 25°C.

The following *Drosophila* stocks were used: i) *w*^*1118*^; *FRT82B, ii) Df(3R)Exel6169,P{XP-U}Exel6169/TM6B,Tb* (#7648) *iii) UAS-RasV12, FRT82B (RAS* ^Hi^ – modified from stock #4847) and iv) *UAS-Ampk*^*Trip20(RNAi)*^ (#57785) - all provided by the Bloomington *Drosophila* Stock Center). *UAS-RasV12; FRT82B (RAS* ^Lo^)^14^ was provided by Tian Xu. *w*^*1118*^ and *Viking*-*GFP* ^53^ were gifts from K. Moberg (Emory University). *Lkb1*^*4A4-2*^ and *Lkb1*^*4B1-11*^ were gifts from J. McDonald (Kansas State University). *Lkb1*^*X5*^ was a gift from W. Du (University of Chicago). Fluorescently labeled mitotic clones were induced in larval eye-imaginal discs using the following strain: *y,w, eyFLP1; Act >y+> Gal4, UAS-GFP (or RFP); FRT82B, Tub-Gal80* (provided by Tian Xu).

### Generation of *Drosophila* Lkb1 antibody

ProteinTech was used to generate a custom Lkb1 polyclonal antibody specific to *Drosophila* using the following peptide sequence: VEDEMTVLLANKNFHYDV-Cys. Guinea Pigs were immunized and supplemented with booster immunizations before final antibody production after 102 days. Antibodies were affinity purified with Elisa confirmation of purification, and final antibody concentrations were estimated by SDS-PAGE.

### BrdU Staining

3^rd^ instar larval eye-imaginal discs were dissected in Grace’s Insect Medium (ThermoFisher) then transferred into Grace’s Insect Medium containing 0.25mg/ml BrdU (Invitrogen B23151) and incubated at 25°C for 90 minutes. Discs were then washed in Grace’s Insect Medium for five minutes on ice followed by washing two times for five minutes each in 1X PBS on ice. Discs were fixed overnight (wrapped in foil) in 1% paraformaldehyde/0.05% Tween20. The following day discs were washed three times for five minutes each in 1X PBS and permeabilized for 20 minutes at RT in 0.3% PBST. To remove detergent, discs were washed five times for five minutes each in 1X PBS and DNAse treated for 30 minutes at 37°C. Discs were then washed three times for 10 minutes each in 0.1% PBST and incubated overnight at 4°C in mouse anti-BrdU primary antibody (B44) (BD, 1:50). The next day, discs were washed 5 times for a total of 30 minutes with 0.1% PBST and incubated overnight in goat anti-mouse F(ab)’2 AlexaFluor-555 secondary antibody (Cell Signaling, 1:500). Finally, discs were washed three times for 10 minutes each in 0.1% PBST and mounted in VectaShield anti-fade mounting medium.

### Cell-cycle analysis

Live GFP-labelled 3^rd^ instar eye-imaginal disc cells were dissected in 1X PBS and simultaneously dissociated with gentle agitation and stained (wrapped in foil) with Hoechst 33342 (Cell Signaling, 500 g/ml) for two hours using a solution of 450µl 10X Trypsin-EDTA (Sigma), 50µl 10X PBS, and 0.5µl Hoechst 33342. Cells were then passed through a 40μm cell strainer prior to FACS analysis. Hoechst 33342 expression was analyzed for a minimum of 10,000 GFP-positive cells by flow cytometry on a Becton Dickinson FACS Canto II cytometer using FACSDiva software. Elimination of dead cells and the distribution of cells within G1, S, and G2/M phases of the cell cycle was determined using FlowJo software.

### Western blotting

Twenty 3^rd^ instar larvae were dissected in 1X PBS and eye-imaginal discs were transferred to a 1.5ml microcentrifuge tube containing 1ml of fresh 1X PBS. Discs were spun down at 4°C for 1 min at 9,600g and supernatant was removed. 2X Laemmli Sample Buffer was added and discs were boiled for 10 minutes at 100°C, and spun down. Approximately 10µg of protein was loaded into a 12% polyacrylamide gel. Alternatively, 3^rd^ instar larvae were dissected and 20µg of crude extract was loaded into a 10% polyacrylamide gel. Samples were run at 100V and separated by SDS-PAGE before transferring to a polyvinylidene difluoride (PVDF) membrane overnight at 0.07amps at 4°C. Membranes were blocked for 1 hour with 10% skim milk in 1X tris-buffered saline plus Tween 20 (TBST) and placed in primary antibody overnight in 1X TBST with 5% skim milk or BSA at 4°C. The following day, membranes were washed three times for 10 minutes each in 1X TBST and placed in secondary antibody in 1X TBST with 5% skim milk or BSA for 1 hour at RT. After three additional 10-minute washes in 1X TBST, ECL-reagent (Amersham, RPN2232) and X-ray film were used to detect signals. When necessary, membranes were stripped using GM Biosciences OneMinute Plus Western Blot Stripping Buffer (GM6011). Primary antibodies and dilution: affinity purified guinea pig anti-*Drosophila* Lkb1 (Protein Tech, 1:1000), rabbit anti-Ras (Cell Signaling 3965, 1:1000), rabbit anti-phospho Ampk (Thr 172) (40HP) (Cell Signaling, 1:1000), mouse anti-*Drosophila* AMPK1/2 (BioRad, 1:1000), rabbit anti-diphosphorylated ERK (Sigma, 1:1000), Rabbit anti-phospho MEK1 (Ser 217+221) (Invitrogen, 1:500), rabbit anti-*Drosophila* phospho p70 S6 Kinase (Thr 398) (Cell Signaling (1:1000), rabbit anti-phospho 4E-BP1 (Thr 37/46) (Cell Signaling, 1:1000), rabbit anti-phospho AKT (Ser 473) (Cell Signaling, 1:1000), mouse anti-phospho CaMKII (Thr 286) (22B1 Santa Cruz Biotechnology, 1:200), rabbit anti-ATG8a (Creative Diagnostics, 0.2g/ml), and mouse anti-actin (JLA20; (Developmental studies Hybridoma Bank, 1:1000).

### Immunostaining

3^rd^ instar larval eye-imaginal discs were dissected in 1X phosphate-buffered saline (PBS) and fixed in 4% paraformaldehyde for 30 minutes on ice. Discs were then washed three times for 10 minutes each in ice cold 1X PBS, permeabilized in 0.3% Triton X100/1X PBS (PBST) for 20 minutes at RT, and washed again three times for 10 minutes each before blocking in 10% normal goat serum in 0.1% PBST for 30 minutes at RT. Discs were incubated in primary antibodies (4°C overnight) in 10% normal goat serum (NGS)/0.1% PBST. The following day, discs were washed three times for five minutes each in 0.1% PBST before incubating in secondary antibodies (in the dark at RT for one hour) in 10% NGS/0.1% PBST. Finally, discs were washed three times for 10 minutes each in 1X PBS at RT and mounted using VectaShield anti-fade mounting medium. Primary antibodies and dilution: rabbit anti-cleaved *Drosophila* DCP1 (Asp216) (Cell Signaling, 1:100), mouse anti-MMP1 (3A6B4/5H7B11/3B8D12 antibodies were mixed in equal amounts) (DSHB, 0.2µg/ml), and Rabbit anti-pAMPK (T172) (Cell Signaling 1:100). Fluorescent secondary antibodies were from Life Technologies. DAPI was used to stain DNA.

### Widefield and confocal imaging

Brightfield adult images were taken using a Leica S6D dissecting microscope. Fluorescent images were taken on a Leica MZ10F (× 1 0.08899 NA) or Leica TCS SP8 inverted confocal microscope (× 10 air HC PL Fluotar, 0.3 NA, × 20 air HC PL APO, 0.75 NA, or × 40 oil HC PL APO, 1.30 NA) using 0.88□μm z-stack intervals and sequential scanning (405□nm DMOD Flexible, 488□nm argon, 514□nm argon). All images were processed using ImageJ/FIJI and compiled in Adobe Photoshop.

### Allografting

Tissue allografting was performed as described previously^54^. 3^rd^ instar larvae were placed in a sterile petri dish containing 1X PBS and washed to remove residual fly food from the larval cuticle. Larval eye-imaginal discs were then dissected in 1X PBS. Sterile forceps were used to mince tissue into small pieces in preparation for implantation. *w*^*1118*^ virgin female host flies were anesthetized with CO_2_ and placed ventral-side up on double-sided sticky tape. Care was used to ensure that flies were well adhered to tape. A 10µl sterile Hamilton Syringe with a 34 gauge 1-inch needle (45° needle angle) was used to aspirate a single piece of eye disc tissue into the needle, loading as little 1X PBS as possible. Forceps were used to hold the host abdomen steady and the syringe needle was used to pierce the abdomen and inject the eye disc tissue. Host flies were then removed from the double-sided tape and moved to a fresh vial of food placed horizontally at all times. Between genotypes the needle was cleaned by pipetting in and out with 1X PBS several times. Flies were monitored daily for survival and GFP-positivity, with transfer to new vials every two days. Death observed during the first 7 days was deemed artefactual, due likely to the injection procedure and not malignant growth. Flies were monitored for a total of 32 days.

### SiMView Light Sheet Microscopy

Prior to mounting, live wandering 3^rd^ instar *Drosophila* larvae and giant larvae (13 days AEL) were selected for stage and proper expression then cooled in a petri dish placed on top of an ice bucket. After sufficiently cooled to minimize movement, the samples were attached posterior side up to a 3mm diameter stainless steel post using gel-control super glue (Ultra Gel Control, Loctite). When mounting, the sample’s mouthparts were adhered in an extended state in order to improve image quality (i.e. reduce object depth) of the tumors. After allowing the adhesive to dry, the sample and post was loaded into an adapter that is magnetically attached to a multi-stage stack with degrees of freedom in the X-Y-Z and rotational directions. The sample chamber is sealed using custom-made rubber gaskets and filled with Schneider’s Medium. The instrument is constructed as previously published with slight modification^55,56^. All data was collected using a Nikon 16x/0.8 NA LWD Plan Fluorite water-dipping objective and Hamamatsu Orca Flash 4.0 v2 sCMOS cameras. Exposure time for all experiments was 15ms per frame. We collected data using a single camera view and two illumination arms, exciting with each arm in sequence for each color and timepoint. In our SIMView implementation for one-photon excitation, multiview image stacks are acquired by quickly moving the specimen over the desired z range and alternating light-sheet activation in the two illumination arms for each volume. This bidirectional illumination and detection capture recordings from two complementary views of each z plane in two illumination steps. Notably, no mechanical rotation of the specimen is required. The switching of laser shutters in the two illumination subsystems is performed within a few milliseconds. GFP and RFP fluorophores were excited using 488nm and 561nm Omicron Sole lasers, respectively.

### Analysis of SiMView Data

Following data acquisition, images were processed prior to analysis. All data had 90 counts subtracted to account for dark counts of the sCMOS cameras. Images from each illumination arm corresponding to the same Z slice were merged and corrected for intensity variation. Details on these algorithms are previously published^57^. Vkg-GFP pixel intensity over time was measured by using maximum intensity projections of 3D volumes from 8 different time points between 0 and 14 hours. The pixel intensity for the tracheal region of interest was measured for each time point using FIJI/Image J. 3D volumetric time-lapse data were visualized using Bitplane Imaris 9 (Fig. 4 c-e). Subsets of the entire 2000-3000 timepoint series (∼3 - 5 TBs in size) were selected for 3D inspection and visualization from maximum intensity projection (MIP) images. 3D regions of interest (3D-ROI) were created using Imaris’ intensity-based Surfaces function.

### Pharmacology

Molasses-based food was melted and 10ml of food was aliquoted to vials. While warm, 10µl of H_2_0 or 10µl of 5mM KN-93 (Millipore Sigma, 422711) were added to vials, respectively. Food vials were cooled and allowed to solidify before use. Vials not immediately used were placed at 4°C. *Adult y,w, eyFLP1; Act >y+> Gal4, UAS-GFP; FRT82B, Tub-Gal80* virgin female flies were crossed to *FRT82B* or *UAS-RasV12*^*Hi*^*/Lkb1*^*4A4-2*^ males, respectively. Flies were moved to embryo “egg-laying cups” and allowed to egg-lay onto grape juice agar plates at 25°C. Flies were moved onto fresh agar plates every 24 hours. After each 24hr period, embryos were collected using forceps and placed onto a fresh vial of food. Embryos were placed at 25°C and allowed to hatch. Once of age, 2^nd^-instar larvae were collected and placed onto drug containing media at 25°C. Survival was quantified as the percentage of total embryos placed that survived to pupation and adulthood.

### Survival analysis of patient data

cBioPortal was used to obtain survival, copy number, mRNA expression, and RPPA expression data available through the Cancer Genome Atlas (TCGA). For survival analysis, specific studies used included: TCGA Pan-Lung Cancer study^58^ and TCGA Lung Adenocarcinoma studies (PanCancer Atlas^59^ and Provisional). Out of 1144 total samples, samples with specific KRAS G12C, G12D, or G12V mutations were selected for further analysis (115 samples for mRNA analysis and 76 samples for copy number analysis). Stratification as KRAS^Lo^ or KRAS^Hi^ was based on normalized (Log2) mRNA expression or relative copy number. Patients with KRAS normalized mRNA expression value less than 10.825 were designated as RAS^Lo^, while patients with a normalized mRNA expression value greater than 10.825 were designated as RAS^Hi^. For copy number analysis, diploid patients were designated as KRAS^Diploid^, while patients with KRAS gains and amplifications were designated as KRAS^Gain/Amp^. Of these patients, mono or biallelic loss or predicted loss-of-function mutations in *LKB1* were also obtained. For pAMPK correlation analysis, specific studies used included: TCGA Pan-Lung Cancer study (Nat Genet 2016) and TCGA Lung Adenocarcinoma study (PanCancer Atlas). Samples with specific KRAS G12C, G12D, or G12V mutations as well as RPPA expression data for pAMPK (T172) were selected for further analysis (n = 71).

### Canonical Circuit Activity Analysis

The HiPathia web-application (http://hipathia.babelomics.org) was used to identify differentially expressed (activated or inhibited) pathways. RNA-sequencing raw RSEM count data based on human genome build hg19 was obtained for the TCGA lung adenocarcinoma (LUAD) patients from the Genomic Data Commons (GDG) legacy archive (https://portal.gdc.cancer.gov/legacy-archive). Patients without all data types were excluded. Patients were obtained and grouped based on KRAS status using CBioPortal as previously described and determined to be KRAS/LKB1WT if concomitant somatic mutations in LKB1 were absent, and as KRAS/LKB1Mut if secondary LKB1 mutations were present. Genes with average counts per million (CPM) of greater than 0.1 across all samples were kept. Normalization was done with TMM (trimmed mean of M-values) method and log2 transformed using edgeR package^60^. The normalized expression matrix was then used as input in HiPathia web-application to identify up- or down-regulated pathways between the two groups (mutant vs. wild-type) against all available pathways in HiPathia. Finally, differential gene fold change was estimated using the Limma R package.

### Statistical Analysis

GraphPad Prism 7 and 8 were used to generate *P* values using the two-tailed unpaired Student’s t-test to analyze statistical significance between two conditions in an experiment, ordinary one-way ANOVA with a Tukey’s multiple comparisons test for experiments with three or more comparisons, and Log-rank (Mantel-Cox) test for analysis of survival data. Significance was assigned to *P* values less than 0.05 unless otherwise indicated. For Figure 6e and f, statistical analysis was conducted using RStudio. Data was divided into two groups, LKB1 loss of function (n = 40) and LKB1 wild type (wt) (n= 31). A single outlier sample in the LKB1 mutation category was excluded and calculated z-score for pAMPK and KRAS expression data was used. The correlation between the AMPK and KRAS was conducted and a spearman’s correlation test. Due to the relatively small sample size, a p-value of <= .1 or 10% was considered significant.

## Supplementary Materials

### Materials and Methods

Fig. S1 Blocking cell death with *P35* in *Lkb1* mutant clones does not phenocopy *Ras*^*Lo*^*/Lkb1*^*-/-*^.

Fig. S2 Validation of genetic and pharmacologic agents.

Fig. S3. High level *KRAS* does not result in survival differences in *TP53* mutant lung cancer patients.

**Movie S1**. Long-term imaging of tumor cell invasion in autochthonous *Kras/Lkb1* malignant tumors.

## Author Contributions

M.G.R., B.R., C.S. and E.K. conceived and designed the project. B.R. and C.S. performed the *Drosophila* and molecular biology experiments. R.E.P contributed to visualization and editing of the manuscript, E.K., N.A., J.M.H., T.L.C., and M.G.R. designed and performed the SiMView imaging experiments. W.G., N.A., E.K., and M.G.R. analyzed the SiMView data. B.D., M.R., and B.R. performed the bioinformatic, correlation studies, and statistical analysis using human patient data, and B.R. and M.G.R. wrote the manuscript.

## Acknowledgements

Thank you to T. Xu, J. McDonald, K. Moberg, and D. Lerit for gifts of fly stocks, reagents, and equipment. We acknowledge the Bloomington Drosophila Stock Center, Vienna Drosophila Resource Center, TRiP at Harvard Medical School (NIH/NIGMS R01-GM084947) and the Developmental Studies Hybridoma Bank (DSHB) for providing fly stocks and antibodies. We thank members of the A. Marcus, W. Zhou, K. Moberg, and R. Read laboratories for helpful comments, discussion, and teaching of techniques. This work was supported in part by the Advanced Imaging Center at the Janelia Research Campus of the Howard Hughes Medical Institute. The Advanced Imaging Center is jointly supported by the Gordon and Betty Moore Foundation and Howard Hughes Medical Institute. Research reported in this publication was supported in part by the Winship Biostatistics and Bioinformatics and Emory Integrated Cellular Imaging Shared Resources of Emory University under NIH/NCI award number P30CA138292. Research reported in this publication was supported by the National Cancer Institute of the National Institutes of Health under Award Number P50CA217691, R01CA194027 (MGR) and R01CA201340 (MGR). The content is solely the responsibility of the authors and does not necessarily represent the official views of the National Institutes of Health.

## Supplementary Materials

### Materials and Methods

#### *Drosophila* stocks and maintenance

Flies were grown on a molasses-based food at 25°C. The following *Drosophila* stocks were used: *FRT82B, UAS-P35, UAS-RasV12; FRT82B (KRAS*^*Lo*^)^*14*^, *and UAS-RasV12, FRT82B (KRAS*^*Hi*^) were provided by Bloomington *Drosophila* Stock Center. *w*^*1118*^ was a gift from K. Moberg (Emory University).

#### Widefield and confocal imaging

Brightfield adult images were taken using a Leica S6D dissecting microscope. Fluorescent images were taken on a Leica MZ10F (× 1 0.08899 NA) or Leica TCS SP8 inverted confocal microscope (× 10 air HC PL Fluotar, 0.3 NA, × 20 air HC PL APO, 0.75 NA, or × 40 oil HC PL APO, 1.30 NA) using 0.88 μm z-stack intervals and sequential scanning (405□nm DMOD Flexible, 488□nm argon, 514□nm argon). All images were processed using ImageJ/FIJI and compiled in Adobe Photoshop.

#### Survival analysis of patient data

CBioPortal was used to obtain survival and mRNA expression data available through the Cancer Genome Atlas (TCGA). For survival analysis, specific studies used included: TCGA Pan-Lung Cancer study (Nat Genet 2016) and TCGA Lung Adenocarcinoma study (PanCancer Atlas). Out of 1144 total samples, samples with specific KRAS G12C, G12D, or G12V mutations were selected for further analysis (115 samples). Stratification as RAS^Lo^ or RAS^Hi^ was based on normalized (Log2) mRNA expression. Patients with a normalized mRNA expression value less than 10.825 were designated as RAS^Lo^, while patients with a normalized mRNA expression value greater than 10.825 were designated as RAS^Hi^. Of these patients, secondary deletions or loss-of-function mutations in P53 or LKB1 were also obtained.

#### Statistical Analysis

GraphPad Prism 7 was used to generate *P* values using the ordinary one-way ANOVA with a Tukey’s multiple comparisons test for experiments with three or more comparisons and Log-rank (Mantel-Cox) test for analysis of survival data. Significance was assigned to p values<0.05. Error bars represent the mean ± SD.

---

**Supplementary Figure 1.**
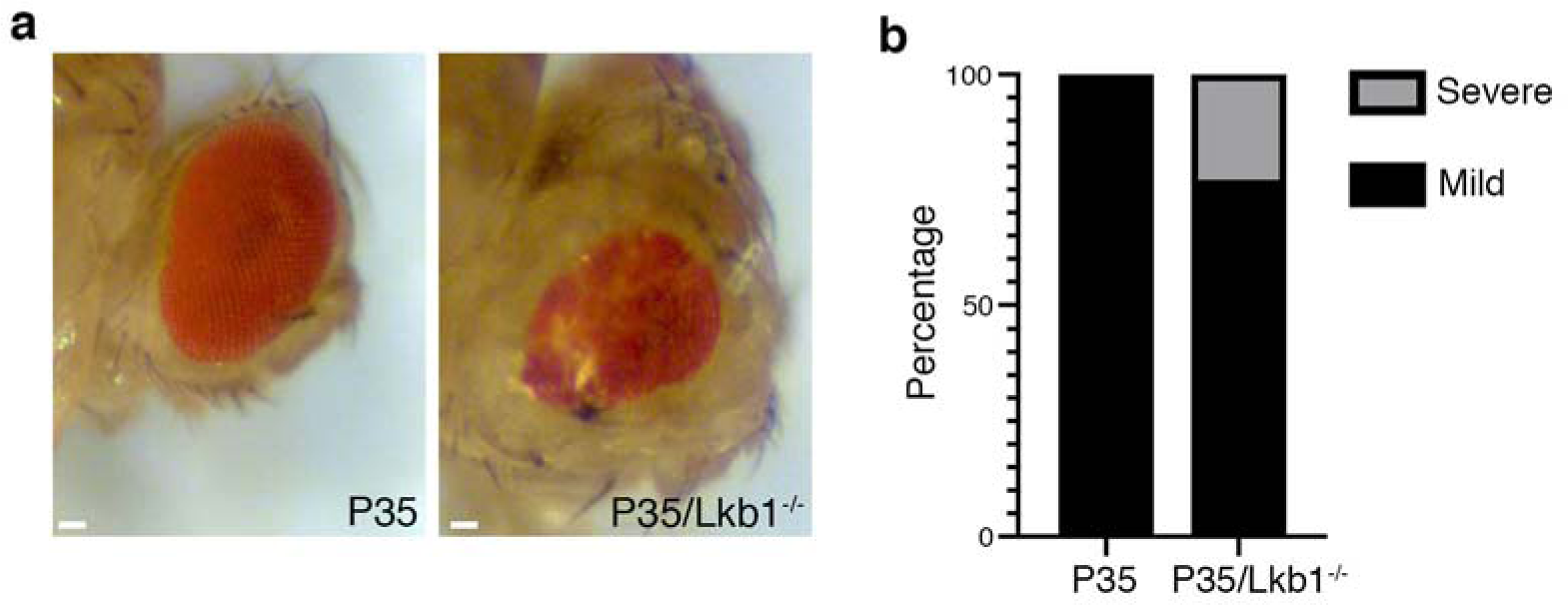
Blocking cell death with *P35* in *Lkb1* mutant clones does not phenocopy *Ras*^*Lo*^*/Lkb1*^*-/-*^. (a) Brightfield images of mosaic adult eyes expressing *P35* (left) or *P35/Lkb1*^*-/-*^ (right). Scale bar, 20µm. (b) Percentage of *P35* or *P35/Lkb1*^*-/-*^ mosaic eyes with either a mild or severe phenotype (severe phenotype is pictured in (a) for *P35/Lkb1*^*-/-*^.

**Supplementary Figure 2.**
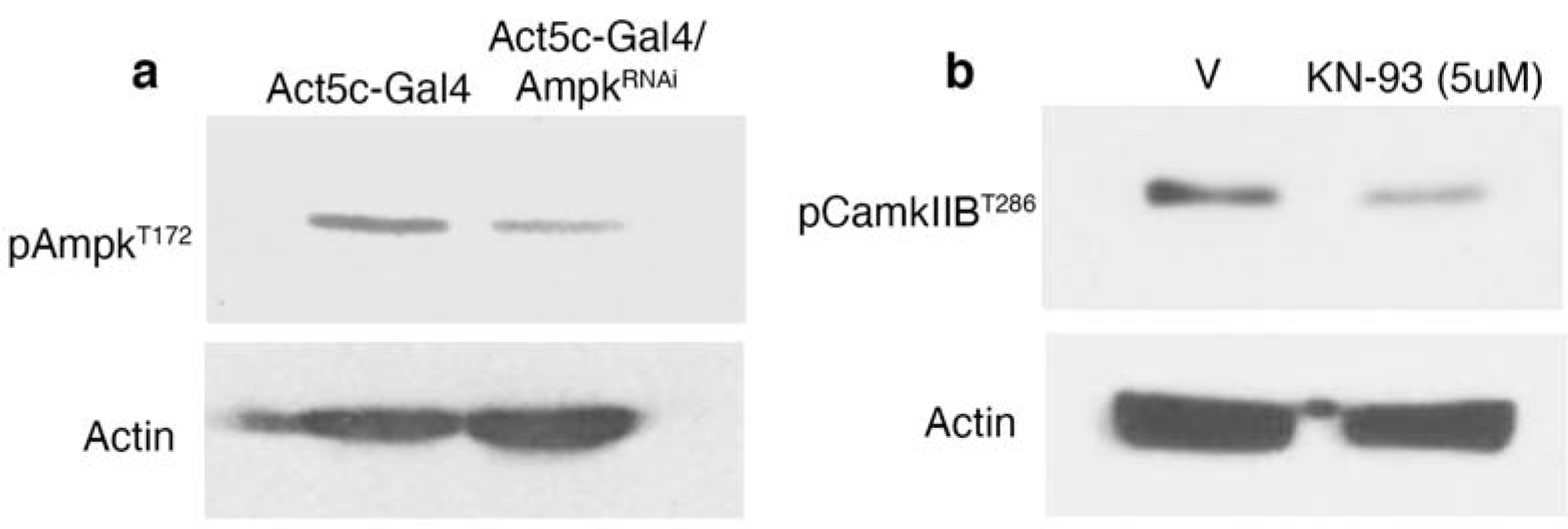
Validation of genetic and pharmacologic reagents. (a) Western analysis of activated Drosophila Ampk from whole larvae of the indicated genotypes. (b) Western analysis of activated Drosophila pCamkIIB from eye/imaginal disc tumors from Ras^Hi^/Lkb1^-/-^ larvae treated with vehicle or KN-93-(5uM).

**Supplementary Figure 3.**
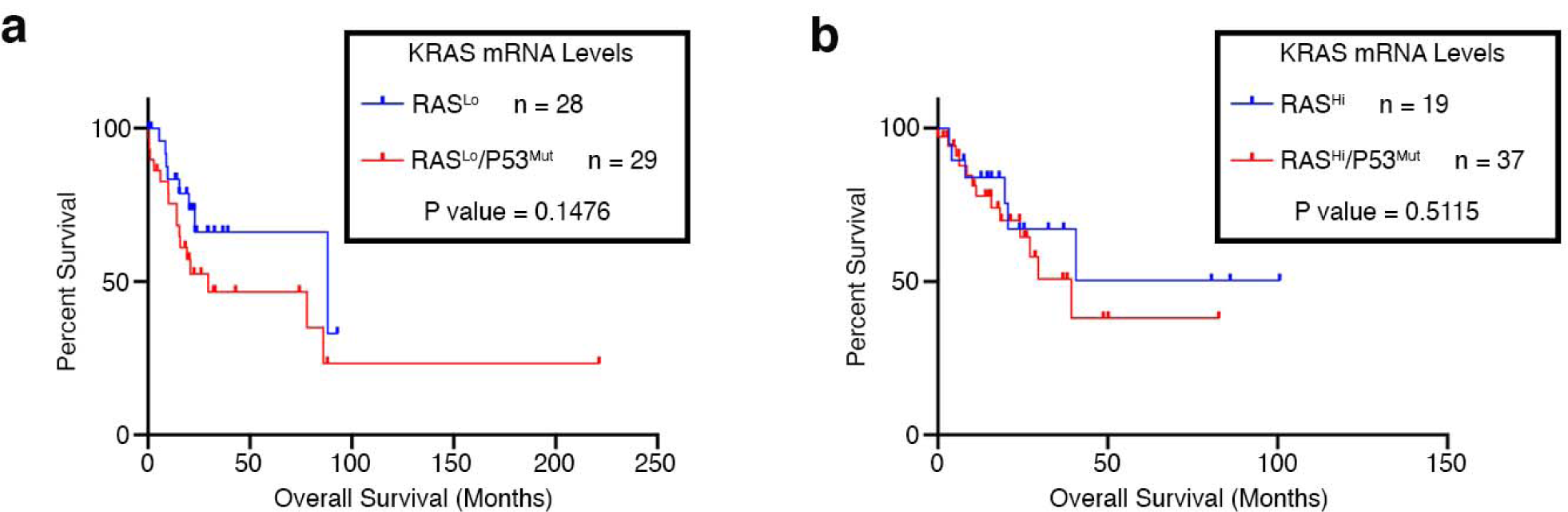
High level *KRAS* does not result in survival differences in *TP53* mutant lung cancer patients. (a-b) Survival analysis using the TCGA Pan Lung Cancer study. Patients were stratified as RAS^Lo^ or RAS^Hi^ using *KRAS* mRNA expression and further stratified based on *TP53* deletion and loss-of-function mutation status. Data were graphed using a Kaplan-Meier survival plot.

